# Single cell transcriptome profiling of mouse and hESC-derived pancreatic progenitors

**DOI:** 10.1101/289470

**Authors:** Nicole A. J. Krentz, Michelle Lee, Eric E. Xu, Shugo Sasaki, Francis C. Lynn

## Abstract

Human embryonic stem cells (hESCs) are a potential unlimited source of insulin-producing β-cells for diabetes treatment. A greater understanding of how β-cells form during embryonic development will improve current hESC differentiation protocols. As β-cells are formed from NEUROG3-expressing endocrine progenitors, this study focused on characterizing the single-cell transcriptomes of mouse and hESC-derived endocrine progenitors. To do this, 7,223 E15.5 and 6,852 E18.5 single cells were isolated from *Neurog3-Cre; Rosa26^mT/^*^mG^ embryos, allowing for enrichment of endocrine progenitors (yellow; tdTomato + EGFP) and endocrine cells (green; EGFP). From a *NEUROG3-2A-eGFP* CyT49 hESC reporter line (N5-5), 4,497 hESC-derived endocrine progenitor cells were sequenced. Differential expression analysis reveals enrichment of markers that are consistent with progenitor, endocrine, or novel cell-state populations. This study characterizes the single-cell transcriptomes of mouse and hESC-derived endocrine progenitors and serves as a resource (https://lynnlab.shinyapps.io/embryonic_pancreas/) for improving the formation of functional β-like cells from hESCs.

## Introduction

Diabetes mellitus is a metabolic syndrome characterized by elevated blood glucose levels that result from reductions in insulin production or action. Insulin is produced by pancreatic β-cells found within the endocrine islets of Langerhans. A potential treatment for diabetes is to replace insulin by transplantation of human embryonic stem cell (hESC)-derived β-cells. Derivation of functional β-cells requires an in-depth understanding of how endocrine cells form during embryonic development.

During mouse and human pancreas development, pancreatic progenitors become restricted to the endocrine cell fate before differentiating to hormone producing cells. This process involves many transcription factors (TFs) that drive the changes in gene expression necessary for endocrine cell genesis. Genetic loss-of-function mouse studies have found a role of individual TFs in the formation of specific islet cell types. From this work, a map of the TF cascade that regulates the formation of endocrine cells, including the β-cells, has emerged (1). However, our understanding of fate decisions during endocrine cell formation is based on studies looking at the whole population of progenitors, using technologies such as bulk RNA-sequencing and often only in mouse cells. The gene expression of individual human and mouse cells during terminal differentiation is unknown.

A promising method to understand gene expression changes at single cell resolution is single cell RNA-sequencing (scRNA-seq). Following the first publication in 2009 (2), commercial platforms and lower sequencing costs have made scRNA-seq a feasible technology for many biologists. Recently, several studies have investigated the single cell transcriptome of healthy and T2D human islets (3–11). From these studies, we have begun to appreciate the cell-type specific gene expression changes that occur during diabetes progression, the differences between mouse and human islets, and the identity of novel islet and pancreatic cell types.

Additionally, two recent studies have begun characterization of the single cell transcriptome of mouse and human progenitors during embryonic development. The first investigated the single cell gene expression of E13.5 embryonic pancreatic cells but very few endocrine progenitors were sequenced (12). The second performed single cell qPCR on 500 cells during several stages of hESC differentiation towards β-like cells (13). In this manuscript, scRNA-seq was used to analyze significantly greater numbers of cells including: 7,223 E15.5 pancreatic cells, 6,852 E18.5 pancreatic cells and 4,497 hESC-derived endocrine progenitor cells. From these data, novel cell types were identified and comparisons between hESC-derived endocrine cells and mouse endocrine progenitors were made. Characterization of these populations will aid efforts to generate an unlimited source of insulin-producing β-cells for diabetes treatment.

## Results

### Strategy for generating single cell transcriptomes of embryonic mouse pancreas

To isolate progenitor populations during mouse embryogenesis, two mouse lines were used: *Neurog3-Cre* and *Rosa26^mTmG^* (Figure 1A). In *Neurog3-Cre; Rosa26^mTmG^* embryos, all cells are labelled with a membrane-targeted Tomato red fluorescent protein (tdTomato). Upon activation of the *Neurog3* promoter, Cre recombinase removes the floxed *tdTomato* cassette, resulting in expression of a membrane-targeted enhanced green fluorescent protein (eGFP). Therefore, cells that have recently activated *Neurog3* express both tdTomato and eGFP marking these cells yellow (EP), while cells that are further along the endocrine cell lineage will express eGFP only (Figure 1A; E) (14). This strategy was used to FACS isolate the three populations from the pancreas of one E15.5 and E18.5 embryo and single cell libraries were generated using 10x Genomics Chromium™ Single Cell 3’ Kit. In total, 7,482 E15.5 and 7,012 E18.5 single cells were sequenced at a depth of >50,000 reads per cell using Illumina NextSeq500.

**Figure 1:**
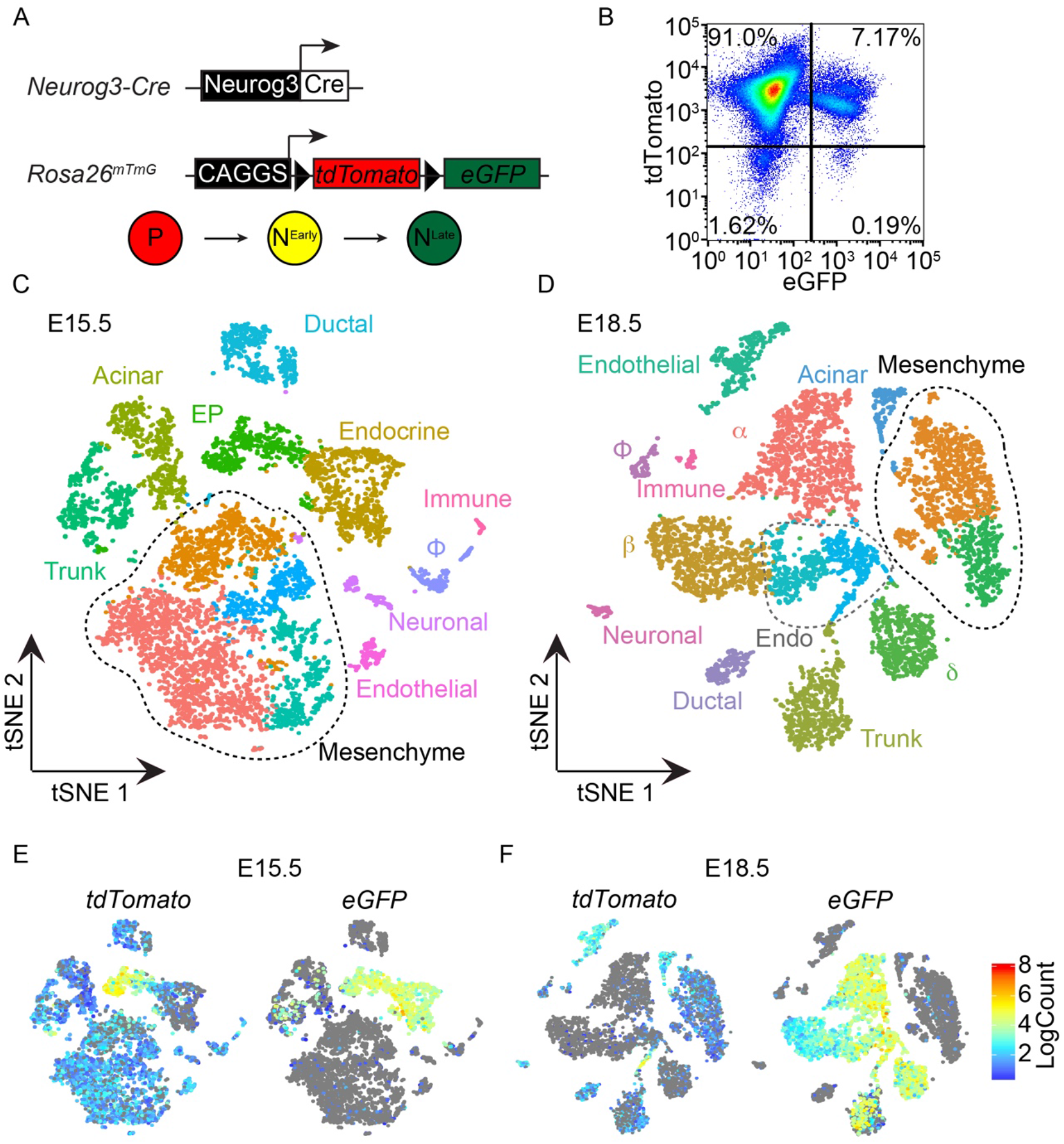
Cell populations in E15.5 and E18.5 mouse pancreas. (A) Schematic overview of the two mouse lines used to isolate cell populations during pancreas development. Using this strategy, pancreatic progenitors (P; red) are tdTomato+, early Neurog3-lineage cells (N^Early^; yellow) are tdTomato+ and eGFP+, and later Neurog3-lineage cells (N^Late^; green) are eGFP+. (B) FACS plot of E15.5 cells used for library generation. (C) Within the E15.5 pancreatic cells there were 13 clusters of nine cell types: trunk, acinar, endocrine progenitor (EP), ductal, endocrine, macrophage (ϕ), neuron, vasculature, and mesenchyme. (D) Within the E18.5 pancreatic cells there were 14 clusters of 11 cell types: trunk, acinar, ductal, maturing endocrine cells (Endo), α-, β-, δ-cells, macrophage (ϕ), neuron, vasculature, and mesenchyme. (E-F) Single cell gene expression of *tdTomato* and *eGFP* at (E) E15.5 and (F) E18.5.

### Identification of cell types in E15.5 and E18.5 pancreas

At E15.5, most cells expressed tdTomato protein only (91.0%). As the yellow (EP; 7.17%) and green populations (0.19%) were less abundant, these cells were pooled together and sequenced as one library (Figure 1B). To explore the cell types present within the E15.5 pancreas, the sequenced red, yellow, and green cells were aggregated into a single dataset using cellranger aggr and low quality cells were excluded from analysis using the R pipelines Scater and Seurat (Materials and Methods). Following this, 7,223 cells were clustered using unsupervised *k*-means clustering and visualized using t-distributed stochastic neighbor embedding (t-SNE) (15), grouping the cells into 13 individual clusters (Figure 1C). Using the top ten genes that are specific for each cluster (Supplementary Table 1), the identity of the cells was inferred. Acinar cells (8.6%) expressed both *Cpa1* and *Cpa2* and duct cells expressed *Krt19* (6.8%), the gene that encodes for CK19 (Figure 1C). Differential expression analysis of the bipotent trunk progenitor cells (7.8%) revealed enrichment for trunk progenitor marker *Sox9* (16, 17), *Spp1* (18), *Mt1*, and *Mt2* (Figure 1C & Figure S1). The endocrine progenitors (EP; 8.3%) expressed *Neurog3, Cdkn1a, Nkx2-2*, and *Pax4* (Figure S1) and were closely associated with endocrine cells (11.4%), which expressed many hormones including *Gcg, Ins1, Ins2, Iapp, Pyy*, and *Gast* (Figure 1C & Figure S1). In addition, there were several rare populations of cells including CD45+ (*Ptprc*) and F4/80+ (*Emr1*) macrophages (ϕ; 2.4%), neurons (2.3%), and endothelial cells (1.9%) (Figure 1C). Finally, most the sequenced cells at E15.5 were found in four clusters of pancreatic mesenchyme cells (50.0%) (Figure 1C).

Next, E18.5 red, green, and yellow single cell libraries were aggregated using cellranger aggr. After filtering, 6,852 cells were clustered using unsupervised *k*-means clustering and the identity of the population was inferred based on the top ten differentially expressed genes (Supplementary Table S2). The most abundant population consisted of pancreatic mesenchyme cells (22.5%) that segregated into two distinct clusters of cells (Figure 1D). There were three endocrine lineages that were named based on gene expression, including alpha-cells (α; 17.8%), the beta-cells (β; 14%), and delta-cells (δ; 8.8%) (Figure 1D). The bipotent trunk progenitor cells (9.6%) continued to express *Sox9*, *Spp1*, *Mt1* and *Mt2* (Figure 1D & Figure S1). There were also two immature endocrine cell clusters (Endo; 11.1%) (Figure 1D). While these clusters shared similar gene expression profiles, one (light blue) expressed endocrine progenitor markers *Neurog3* and *Neurod1*, while the other cluster (teal) expressed β-cell markers *Ins1*, *Ins2* and *Iapp* (Figure 1D & Figure S1): suggesting differences in differentiation or lineage. As was seen in the E15.5 embryo, there were endothelial (6.2%), acinar (3.2%), ductal (3.0%), macrophage (ϕ; 1.8%), and neuronal cells (1%) (Figure 1D).

To verify the library identity of each individual cell, the GRCm38 genome used for alignment was annotated to include the sequences for the *tdTomato* and *eGFP* transgenes. At E15.5 and E18.5, the trunk, acinar, ductal, mesenchymal, endothelial, neuronal, and macrophage cells expressed *tdTomato*, consistent with the non-endocrine lineage of these cell types (Figure 1E; F). At E15.5, the endocrine progenitors (EP) co-expressed both *tdTomato* and *eGFP* (Figure 1E), suggesting recent activation of *Neurog3*. In addition, a subset of immature endocrine cells (endo) at E18.5 co-expressed *tdTomato* and *eGFP*, suggesting that these cells were endocrine progenitors (Figure 1F). As Neurog3+ cells arise from bipotent trunk epithelial progenitor cells, a subset of the trunk cells expressed *eGFP* at both E15.5 and E18.5 (Figure 1E; F). Finally, all endocrine cells at E15.5 and E18.5 expressed *eGFP* (Figure 1E; F), consistent with their derivation from a Neurog3+ endocrine progenitor.

### Characterization of the endocrine cell transcriptome in E15.5 pancreas

To understand the transcriptional changes that occur during endocrine specification, the yellow and green cells were further characterized at E15.5. After filtering, 1350 cells were analyzed using unsupervised *k*-means clustering and visualized using a t-SNE plot (Figure 2A). Seven clusters representing several cell populations were identified using the top ten expressed genes (Supplementary Table 3): endocrine progenitors (EP; 28.7%), alpha-cells (α; 26.4%), β-cells (β; 18.4%), *Chga*-expressing immature endocrine cells (C; 16.7%), trunk progenitor cells (T; 4.7%), ghrelin-cells (ε; 3.9%), and macrophages (ϕ; 1.2%). To confirm the identity of these cells, the expression of several genes was investigated across cell clusters. *Neurog3* was highly expressed in the EP cluster and in a subset of ghrelin cells, while low level expression was also found in *Chga*-expressing cells and trunk (Figure 2B & Figure S2B). *Neurod1*, a target of Neurog3, was expressed throughout the endocrine cell lineage (Figure S2B). The immature endocrine cell cluster expressed both *Chga* and *Chgb* (Figure S2C; green). Both *Ins1* and *Ins2* were expressed in the β-cells while *Gcg* and *Ghrl* were specific to the alpha- and ghrelin-cells, respectively (Figure 2B & Figure S2B).

**Figure 2:**
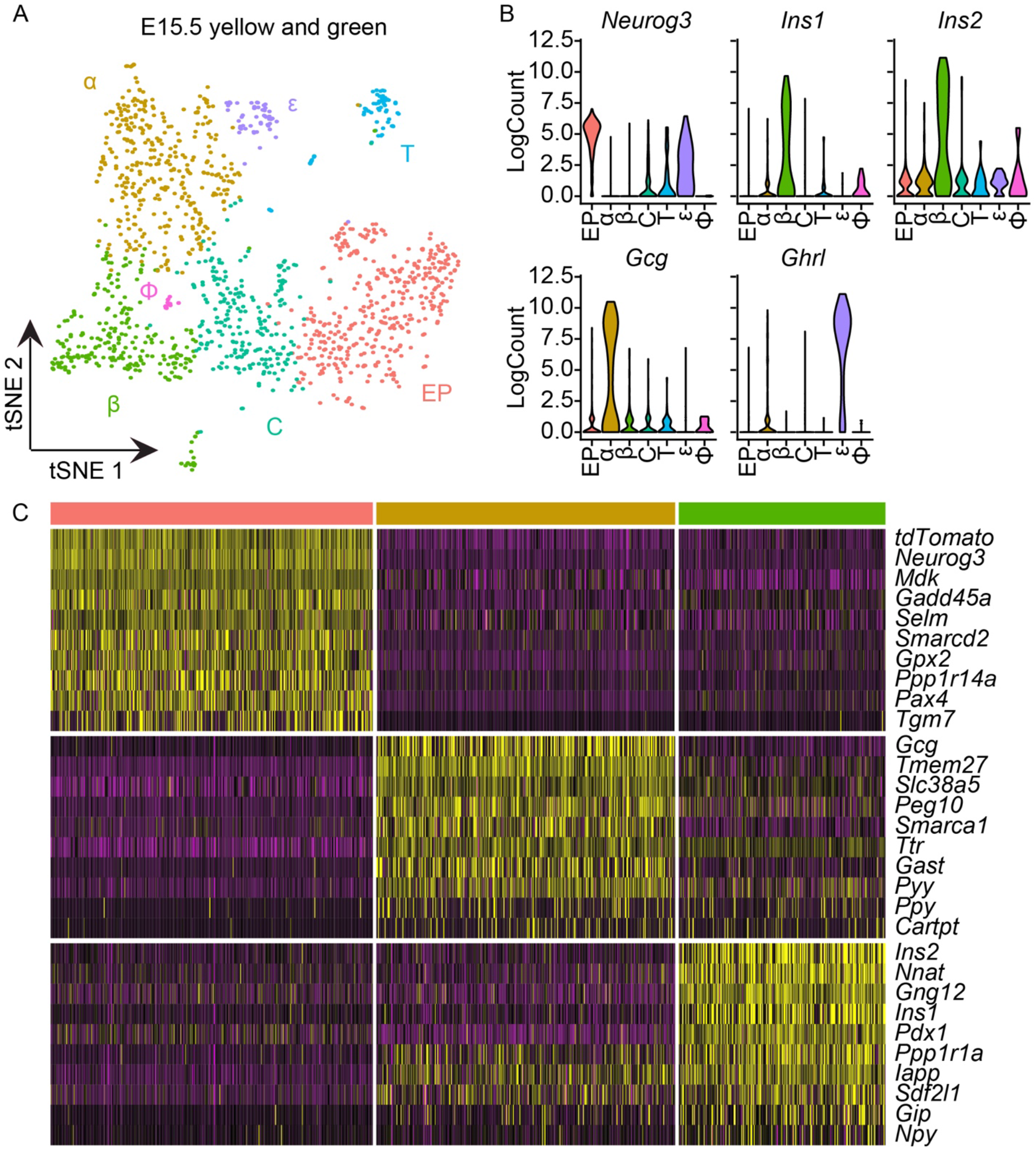
Cell populations in mouse E15.5 yellow and green cells. (A) Clustering of E15.5 yellow and green cells revealed seven clusters. These clusters were identified as endocrine progenitors (EP; 28.7%), *Chga*-expressing immature endocrine cells (C;16.7%), alpha-cells (α; 26.4%), β-cells (β; 18.4%), ghrelin-cells (ε; 3.9%), trunk cells (T; 4.7%), and macrophages (ϕ; 1.2%). (B) Single cell gene expression of *Neurog3*, *Ins1*, *Ins2*, *Gcg*, and *Ghrl* across cell clusters. (C) Heat map of the top ten differentially expressed genes in EP (pink), α-cells (yellow), and β-cells (green) populations.

To find cell-type specific markers, the top ten highly expressed genes in the EP (pink), α-(yellow) and β-cells (green) cells were profiled (Figure 2C). The EP were enriched for expression of known marker genes such as *Neurog3*, and *Pax4* along with novel genes, including *Midkine* (*Mdk*) and *Growth Arrest and DNA Damage Inducible Alpha* (*Gadd45a*) (Figure 2C). The alpha-cells were enriched for expression of *Gcg* and previously proposed alpha-cell markers *Slc38a5* (12) and *Transthyretin* (*Ttr)* (19). The β-cell cluster expressed several β-cell genes, including *Ins1, Ins2, Pdx1*, and *Iapp* (Figure 2C & Figure S2B), along with Neurod1 target *Nnat* (20) and previously identified β-cell marker *Ppp1r1a* (21). Together, these results highlight the utility of single cell transcriptomics to identify novel markers of cell types.

As Neurog3+ progenitor cells exit the cell cycle during differentiation to endocrine cells (22–24), the cell cycle stage of individual cells at E15.5 was investigated. While the EP cluster included dividing cells, cells of the endocrine lineage mainly expressed G1 markers, consistent with cell cycle exit (Figure S2A). Interesting, the trunk population of cells also contained many S- and G2/M-phase cells and had a similar gene expression profile to the trunk population of cells in E15.5 aggr: *Spp1*, *Mt1*, and *Mt2* (Figure S2C; blue). The expression of *tdTomato, eGFP*, and *Neurog3* (Figure S2B; D) suggests that the trunk cluster of cells were bipotent progenitor cells that recently activated the *Neurog3* promoter.

### Pseudotime analysis of E15.5 pancreatic cells

Next, Monocle was used to order the E15.5 library in pseudotime (25–27). For these analyses, the cells previously named as duct, acinar, trunk, EP and endocrine were used to construct a minimum spanning tree based on their transcriptional similarities. This unsupervised algorithm generates a differentiation ‘trajectory’ in pseudotime that models the progression a progenitor cell makes during cell fate decisions (Figure 3A). The three branches represent the terminal differentiation cell types of duct (yellow), acinar (red), and endocrine (green) with a cell fate decision point localized in the center. Along the endocrine lineage, the cells progress through a trunk (purple) and EP (blue) cell fate before becoming endocrine cells (Figure 3A). Pseudotime ordering suggests that ductal cells form first from trunk progenitors while the endocrine cell fate forms later in pseudotime (Figure 3B).

**Figure 3:**
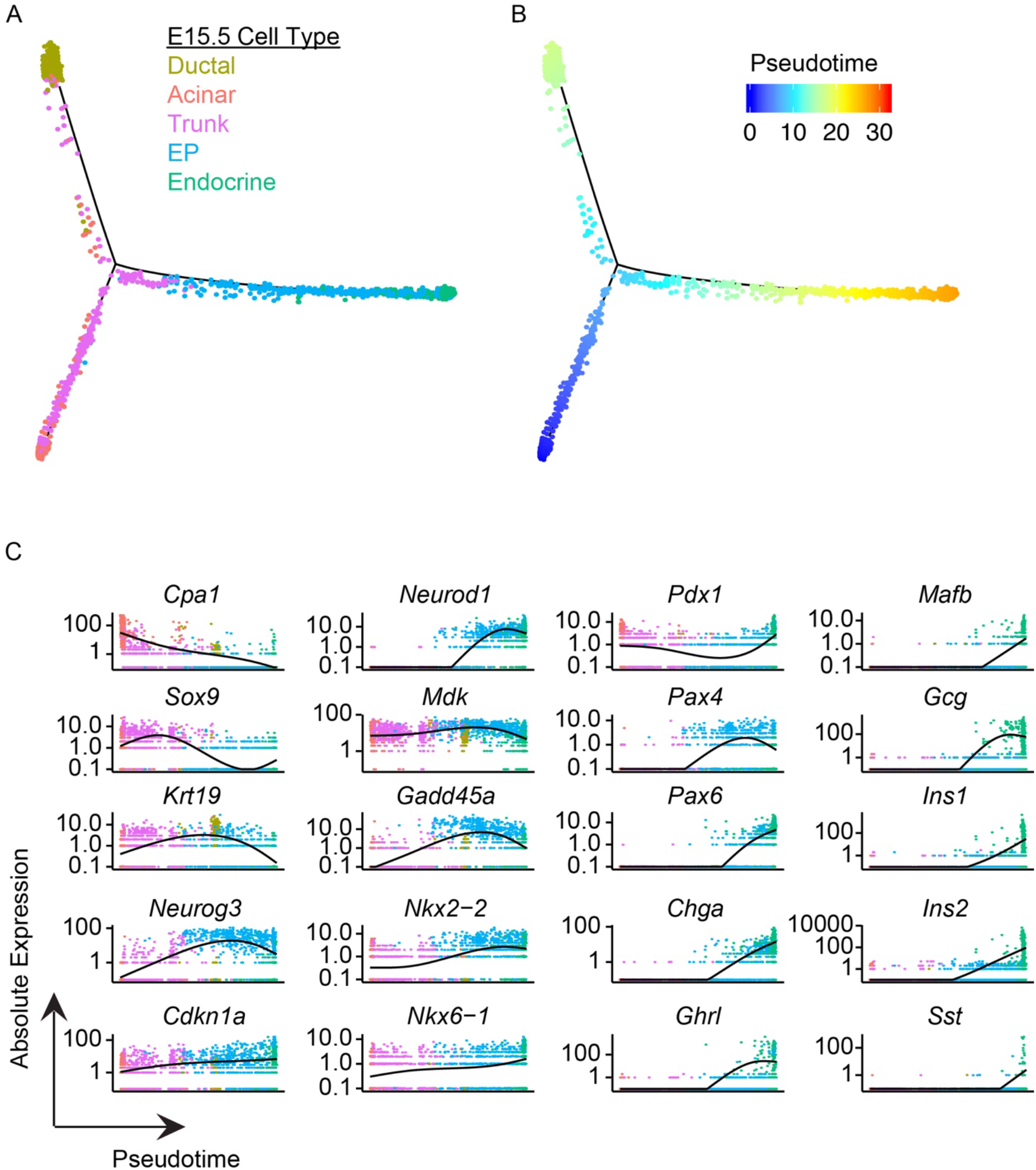
Pseudotime analysis of E15.5 pancreatic cells. (A) Minimal spanning tree for E15.5 ductal (yellow), acinar (red), trunk (purple), endocrine progenitor (EP; blue), and endocrine cells (green). For this analysis, the top 1000 highly variant genes were determined using Seurat and cell states were ordered by Monocle. (B) Pseudotime from 0 (blue) to 30 (red) orders trunk cells first, following by ductal and then the endocrine lineage. (C) Gene expression of cell-type specific markers during pseudotime: acinar markers *Cpa1*; ductal markers *Sox9* and *Krt19*; endocrine progenitor markers *Neurog3*, *Cdkn1a*, *Neurod1*, *Mdk*, and *Gadd45a;* pancreatic markers *Nkx2-2, Nkx6-1*, *Pdx1, Pax4*, and *Pax6*; and endocrine markers *Chga*, *Ghrl*, *Mafb*, *Gcg*, *Ins1*, *Ins2*, and *Sst*.

The trunk progenitor cells were found along all three branches, suggesting that trunk cells are already fated towards the endocrine, ductal, or acinar lineage (Figure 3A). To understand how these cells differ, a differential gene expression analysis was performed on the three populations of trunk cells using Seurat. Within the acinar fated trunk cells, both *Cpa1* and *Cpa2* were among the top ten differentially expressed genes (Figure S3A). In addition, several cell cycle-dependent genes, such as *Top2a* and *Hist1h2ap*, were upregulated in the acinar lineage, suggesting that the trunk cells fated to acinar lineage are actively cycling (Figure S3A). The number of trunk progenitors that fell along the ductal lineage was the smallest and included ECM genes, *Col3a1*, *Col1a1*, and *Col1a2* (Figure S3A). Finally, the trunk cells that were part of the endocrine lineage upregulated expected endocrine cell genes, *Neurog3* and *Cdkn1a* (Figure S3A).

To understand the transcriptional changes that occur during differentiation, we next analyzed the expression of several key genes involved in acinar, ductal and endocrine cell fates. The acinar marker *Cpa1* was highly expressed early in the acinar population before declining over pseudotime (Figure 3C). Next, *Sox9* was upregulated in the trunk progenitor cells before *Krt19* transcription was activated in ductal cells (Figure 3C). As pseudotime continues to the EP population, *Neurog3* was upregulated following by a slow increase in the cell cycle inhibitor *Cdkn1a* (Figure 3C). Expression of *Mdk* and *Gadd45a* mirrored the increase in expression of *Neurog3*, suggesting that these two genes may be involved in endocrine cell formation (Figure 3C). There was a gradual increase in *Nkx2-2*, *Nkx6-1*, *Pax4* and *Pax6* over time, consistent with their known roles in endocrine cell formation, while *Pdx1* was downregulated in the EP population before upregulation in the endocrine cells (17). Finally, endocrine specific genes begin to increase in the EP population and were highest at the end of pseudotime. The pan-endocrine marker *Chga* increased first followed by the *Ghrl*, *Gcg*, *Ins1*, *Ins2*, *Mafb*, and *Sst* (Figure 3C). The sequential upregulation of these genes is consistent with the developmental order of the formation of endocrine cells (28). Taken together, these data confirm the progression of individual pancreatic cells during endocrine cell differentiation and highlights the utility of Monocle.

### Characterization of endocrine cell population at E18.5

Having examined endocrine cell types at E15.5, we next aimed to characterize cells of the endocrine lineage at E18.5. To do this, cells from the E18 yellow and E18 green libraries were pooled and filtered using Scater and Seurat pipelines, resulting in 4,177 cells made up of 593 yellow and 3,516 green cells (Table 1). Visualizing this data using t-SNE revealed 11 clusters: trunk, EP, three β-cell populations, Ghrl cells, alpha-cells, delta-cells, stellate, S-phase cells, and macrophages (Figure 4A). The trunk, EP, stellate and macrophage cells were found in the E18.5 yellow library and expressed *tdTomato* and *eGFP* (Figure 4B; C). Many of the same genes were expressed in E18.5 trunk cells (*Spp1*, *Mt1*, and *Mt2*) while EP expressed *Neurog3*, *Gadd45a*, *Btg2* and *Pax4* (Figure S4B).

**Figure 4:**
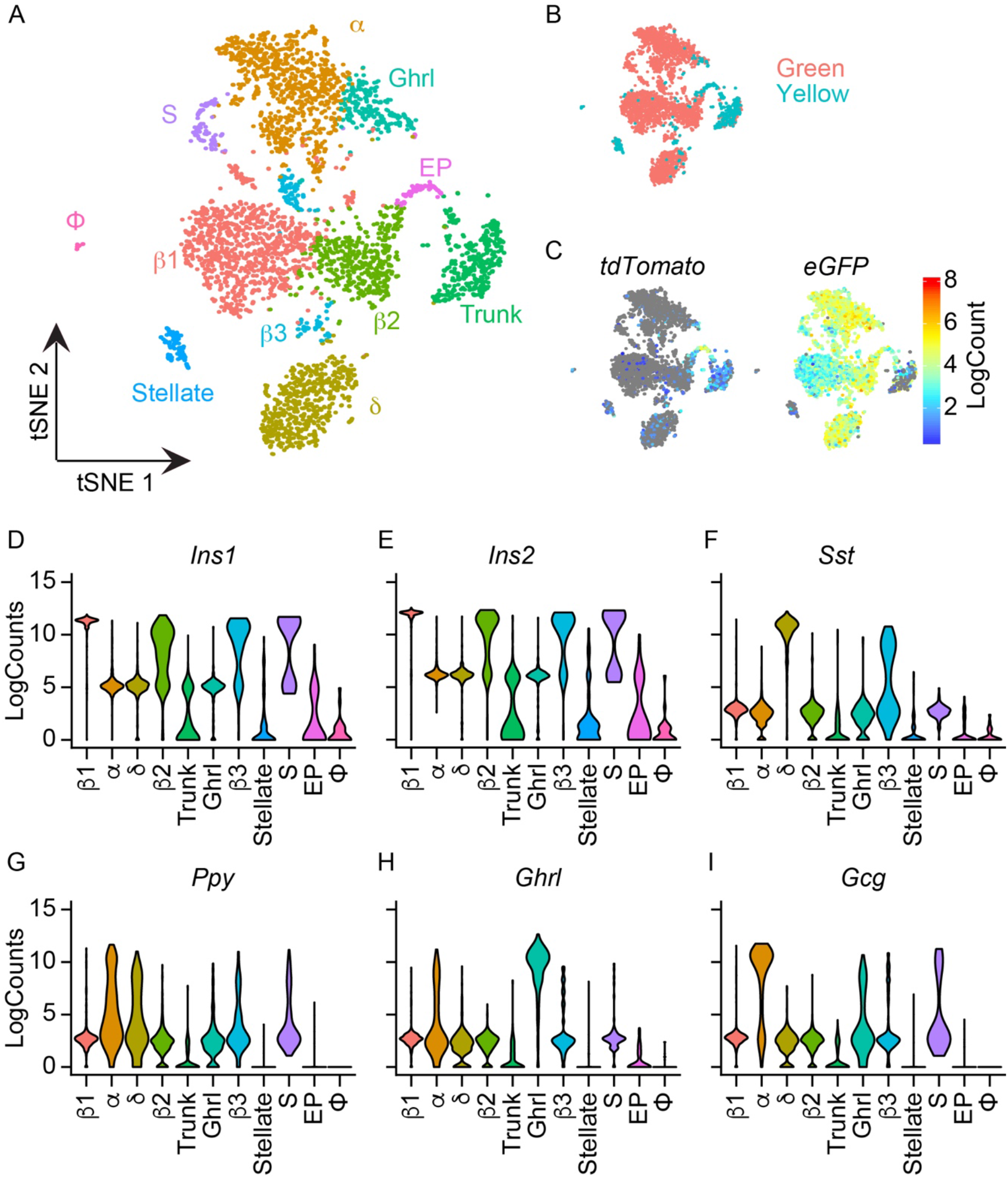
Cell populations in E18.5 yellow and green cells. (A) tSNE plot of 11 cell clusters from E18.5 yellow and green cells: alpha-cell (α), three β-cell (β1, β2, β3), delta-cell (δ), Ghrl-cell, S-phase cells (S), trunk, endocrine progenitor (EP), stellate, and macrophages (ϕ). (B) Library identity of single cells in tSNE plot. (C) Single cell expression of *tdTomato* and *eGFP*. (D-I) Expression of endocrine hormones (D) *Ins1*, (E) *Ins2*, (F) *Sst*, (G) *Ppy*, (H) *Ghrl*, and (I) *Gcg* across clusters.

**Table 1:**
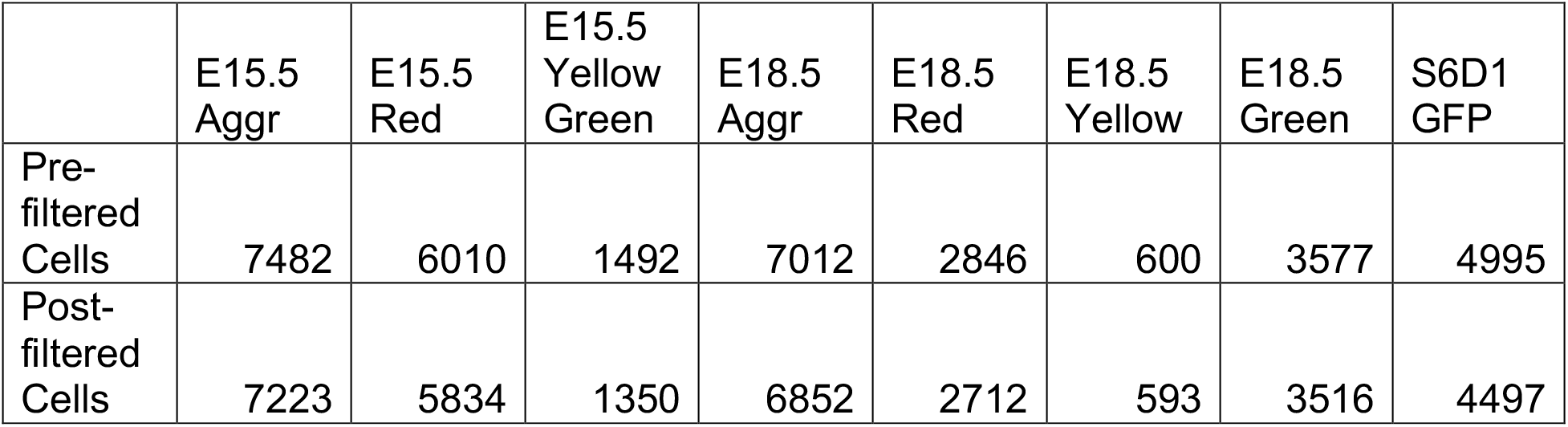
Number of cells pre- and post-filtering for each individual library.

|  | E15.5<br>Aggr | E15.5<br>Red | E15.5<br>Yellow<br>Green | E18.5<br>Aggr | E18.5<br>Red | E18.5<br>Yellow | E18.5<br>Green | S6D1<br>GFP |
| --- | --- | --- | --- | --- | --- | --- | --- | --- |
| Pre-<br>filtered<br>Cells | 7482 | 6010 | 1492 | 7012 | 2846 | 600 | 3577 | 4995 |
| Post-<br>filtered<br>Cells | 7223 | 5834 | 1350 | 6852 | 2712 | 593 | 3516 | 4497 |

The endocrine cells were found within the E18.5 green cell library and expressed only *eGFP* (Figure 4B; C). Both *Ins1* and *Ins2* were highly expressed in the three β-cell populations, β1, β2, and β3, as well as in the S-phase cells (Figure 4D-E), consistent with the start of the wave of replication that is required for β-cell function (29). Differential expression analyses of the β-cell populations reveal cluster-specific differences in gene expression (Supplementary Table S4). The top ten differentially expressed genes in the β1 cluster included maturity markers *Ins1*, *Ins2*, *G6pc2*, and *Slc2a2*, while the β2 cluster was a *Mafb*-expressing immature cell state (Figure S4A). The S-phase cells expressed high levels of *Ins1*, *Ins2*, and *Gcg* suggesting this cluster represents a mixture of alpha- and β-cells (Figure 4D; E; I). Differential gene expression revealed that the S-phase cluster expressed markers of DNA replication, *Top2a* and *Cdk1* (Figure S4A; purple). *Sst* expression was upregulated in the delta cell population and in a subset of the β3 population (Figure 4F). None of the clusters showed specific upregulation of *Ppy* but some alpha, delta and β3 had high *Ppy* expression (Figure 4G). *Ghrl* was upregulated in the Ghrl cluster (Figure 4H) while *Gcg* was highly expressed in the alpha-cell cluster (Figure 4I).

To further investigate the heterogeneity of embryonic endocrine cells, the E18.5 green library was studied. Following filtering, 3,516 cells were visualized using t-SNE plots and the following cell types were identified based on gene expression (Supplementary Table 5): trunk, alpha (α), Ghrl, PP, delta (δ), S-phase cells (S), mitotic cells (M), and two β-cell populations (β1 and β2) (Figure 5A). To confirm these cell identities, the expression of endocrine hormones was determined in single cells. Both *Ins1* and *Ins2* were found in the cells of β1, β2, S, and M clusters, while *Gcg* expression was specific to the α cell populations (Figure S5B). The δ-cell cluster expressed *Sst* and *Hhex* (Figure S5B; C). The Ghrl cells expressed *Ghrl* and PP cells contained *Ppy* transcripts (Figure S5B). Many of the differentially expressed genes for the α-cell lineage were also expressed in the Ghrl and PP clusters (Figure S4C).

**Figure 5:**
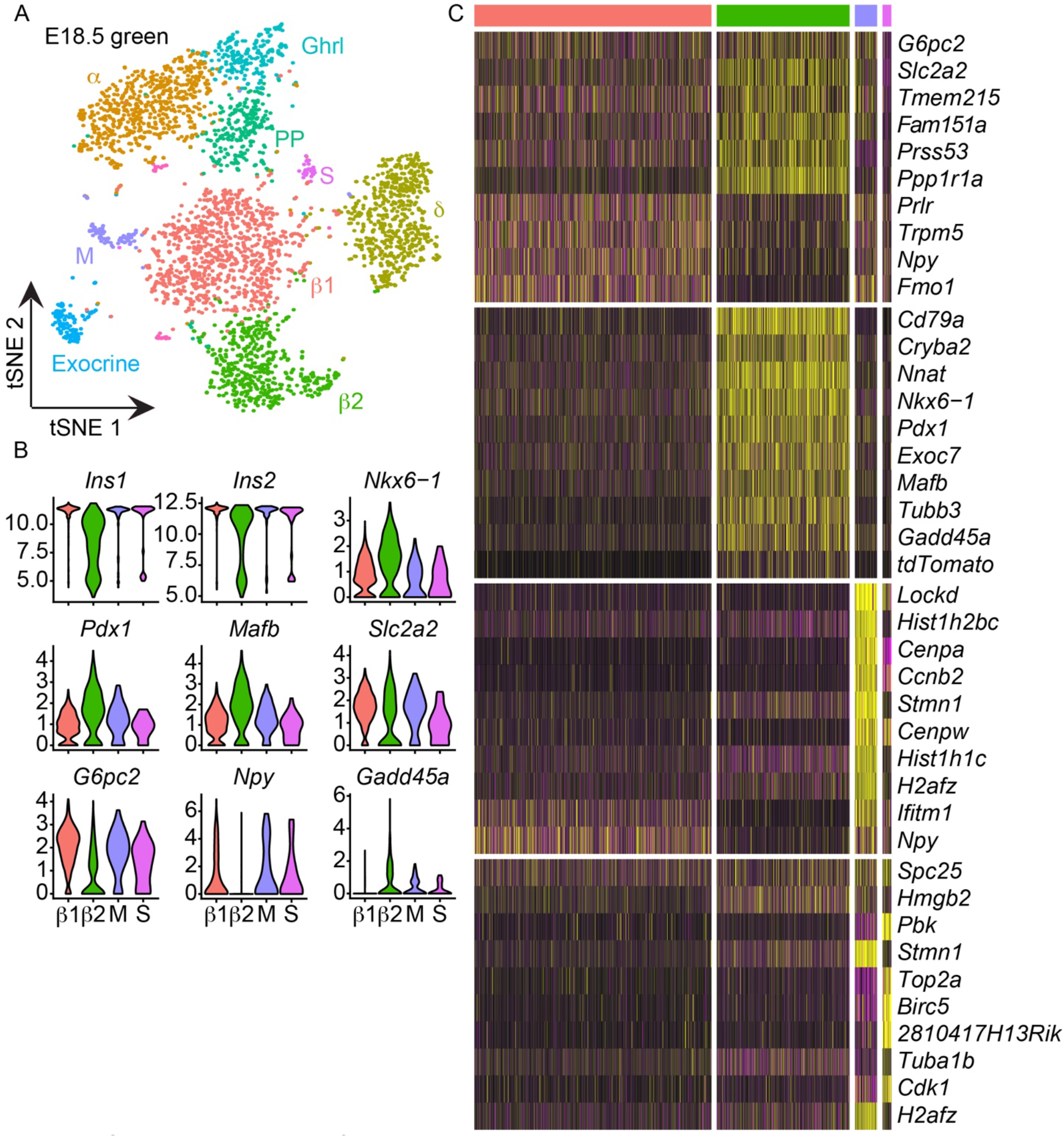
Characterization of endocrine cells in E18.5 green cells. (A) tSNE plot of 10 cell clusters from E18.5 green cells. (B) Violin plots of average gene expression of *Ins1*, *Ins2*, *Nkx6-1*, *Pdx1*, *Mafb*, *Slc2a2*, *G6pc2*, *Npy*, *and Gadd45a* in β1 (red), β2 (green), M (purple), and S (pink). (C) Top ten differentially expressed genes in the β-cell clusters: β1 (red), β2 (green), S (pink), and M (purple).

To understand the heterogeneity within the β-cell populations the expression of the top ten genes for β1 (pink), β2 (green), M (purple), and S cells (pink) was determined (Figure 5C). Most these cells were in the β cluster and expressed genes involved in glucose metabolism including *Slc2a2* and *G6pc2* and expressed high levels of *Ins1* and *Ins2* (Figure 5C). The top ten differentially expressed genes of the β2 cluster included progenitor markers *Pdx1*, *Mafb, Cryba2, Nkx6-1* and *Gadd45a*, suggesting they may be an immature β-cell population (Figure 5C). The other *Ins*-expressing cells are located within the S- and M-phase clusters. The S cluster expressed genes specific to the S phase, such as *Cdk1, Topa2*, suggesting these cells represent a small (1%) population of β-cells undergoing DNA replication (Figure 5C & Figure S5A). In the M cluster (2.5%), the cells expressed the G2/M gene *Ccnb1*, the histone genes *Hist1h2bc*, *Hist1h1c*, *H2af2*, and the kinetochore protein *Spc25*, suggesting that these cells are undergoing mitosis (Figure 5C & Figure S5A). To confirm the identity of these clusters, the cell cycle phase of individual cells in the E18.5 green library was determined using Seurat. The S-phase cluster contained both S- and G2/M-phase cells, while the M cluster had mainly G2/M-phase cells (Figure S5A).

Previously studies in adult β-cells suggest proliferation is accompanied by a decrease in the function and maturation of β-cells (30, 31). To understand if a similar process occurs during mouse β-cell development, we profiled the expression of several β-cell maturity and progenitor markers in the β1, β2, S and M phase populations of cells (Figure 5B). The β2 cluster contained a subset of cells with lower *Ins1*, *Ins2*, *Slc2a2*, *G6pc2*, and *Npy*, consistent with the immature state of these cells (Figure 5B). In addition, this cluster showed an upregulation of genes associated with an immature cell state: *Nkx6-1*, *Pdx1*, *Mafb*, and *Gadd45a* (Figure 5B). Interestingly, the cells of the S and M cluster exhibit a similar gene expression profile as the β1 cluster, suggesting that in the embryonic state proliferation does not reduce maturation (Figure 5B).

### Single cell transcriptome of NEUROG3-lineage during hESC differentiation

To profile the transcriptome of human endocrine progenitors, a CyT49 *NEUROG3-2A-eGFP* hESC reporter line (N5-5) was used (32). N5-5 cells were differentiated using Rezania *et al*. protocol (33) and collected for scRNA-seq before the transition to stage 6 (S6D1) during which the differentiated cells are similar to immature endocrine cells. After filtering, 4,497 GFP+ cells were visualized using t-SNE, revealing nine clusters (Figure 6A). These clusters can be classified as five cell types based on gene expression: endocrine progenitors (EP; 40%), polyhormonal endocrine (Endo; 42.3%), duct (5.7%), liver (8.6%), and an unknown cell type (4.6%) (Figure 6A).

**Figure 6:**
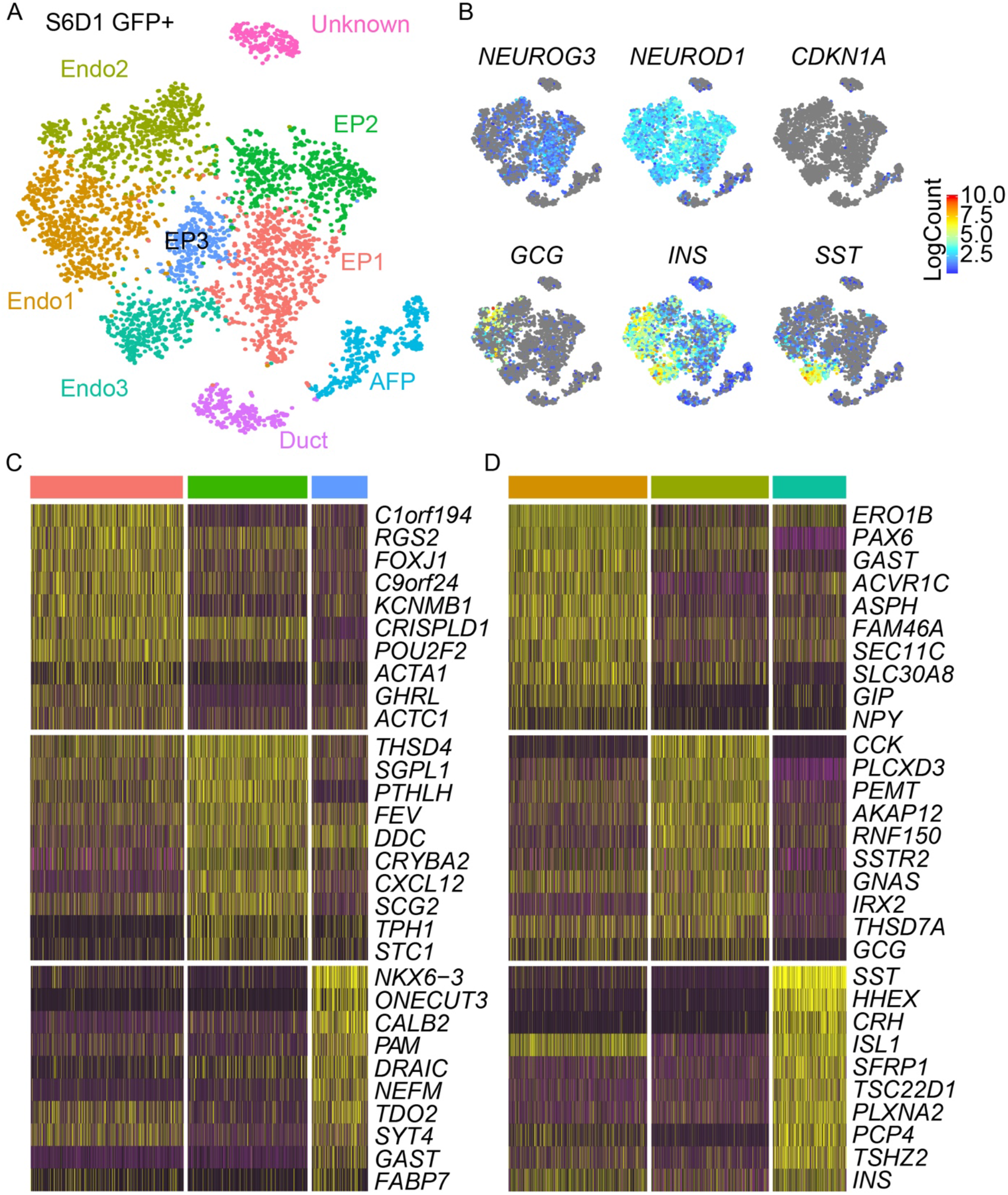
Characterization of *NEUROG3-eGFP* S6D1 cells. (A) tSNE plot of 9 cell clusters from S6D1 GFP+. (B) Single cell gene expression of *NEUROG3*, *NEUROD1*, *CDKN1A*, *GCG*, *INS*, and *SST*. (C) Top ten differentially expressed genes in the endocrine progenitor (EP) cell clusters: EP1 (red), EP2 (green), and EP3 (blue). (D) Top ten differentially expressed genes in endocrine cell clusters: Endo1 (yellow), Endo2 (green), and Endo3 (blue).

Further examination of the expression of endocrine specific genes supported these cell classifications. While all cells expressed GFP protein, only a few cells localized to the EP cluster and expressed *NEUROG3* transcript (Figure 6B). *NEUROD1*, a direct target of NEUROG3 (34), was widely expressed throughout the EP and endocrine clusters (Figure 6B). Interestingly, *CDKN1A*, the downstream target of NEUROG3 that reinforces cell cycle exit during murine endocrine cell differentiation (22), is not abundantly expressed in hESC-derived endocrine progenitors or endocrine cells (Figure 6B). To understand how the three EP clusters differ, the top ten genes that are specific for each individual cluster was determined. The largest EP cluster (EP1) expressed *GHRL*, a gene that marks a multipotent progenitor that can give rise to alpha, PP, and rare β-cells in the mouse (35) (Figure 6C). EP2 contains genes that are associated with serotonin production, including *TPH1* and *FEV* (36) (Figure 6C). In EP3, the *GAST* gene is upregulated, consistent with previous reports of *GAST* induction in INS+ cells during hESC differentiation (37) (Figure 6C).

The endocrine cell population was made up of hormone+ cells, many of which co-expressed multiple hormones including *GCG*, *SST*, and *INS* (Figure 6B). Of the three hormones, *INS* was the most abundantly expressed and can be detected in EP and endocrine cells (Figure 6B). Differential gene expression analysis between the three endocrine clusters revealed an enrichment of β-cell genes *ERO1B* (38), *SLC30A8* (39), and *NPY* (40) in Endo1, suggesting these cells are differentiating β-cells (Figure 6D). The Endo2 cluster appeared fated towards the alpha-cell based on expression of *GCG*, *PEMT* (10), and *IRX2*, while the expression of *SST* and *HHEX* in Endo3 is suggestive of the delta-cell fate (Figure 6D).

### Comparison of mouse and hESC-derived endocrine cells

To understand the “developmental age” of hESC-derived endocrine cells, the single cell transcriptome of S6D1 GFP+ cells was compared to mouse endocrine cells. To do this, all cells of the E15.5 yellow and green library (Figure 2A) and hESC library (Figure 6A) were merged into one dataset, unsupervised clustering was performed, and visualized using a tSNE plot revealing ten clusters (Figure 7A). Using the original identities of the E15.5 yellow and green cells (Figure 2A), the EP, *Chga*-expressing immature endocrine cells, alpha and β-cells clusters were labelled (Figure 7A). These clusters contained a mixture of mouse and human cells, with human cells representing 83%, 70%, 78% and 12% of the Chga, Alpha, Beta, and EP cells, respectively (Figure 7B). While the mouse cells were evenly split among the four clusters, the human cells mainly clustered within the Chga cell type (45%) and very few (1%) were identified as EP cells (Figure 7C). Together, these data suggest that the hESC-derived cells are most like E15.5 *Chga*-expressing immature endocrine cells.

**Figure 7:**
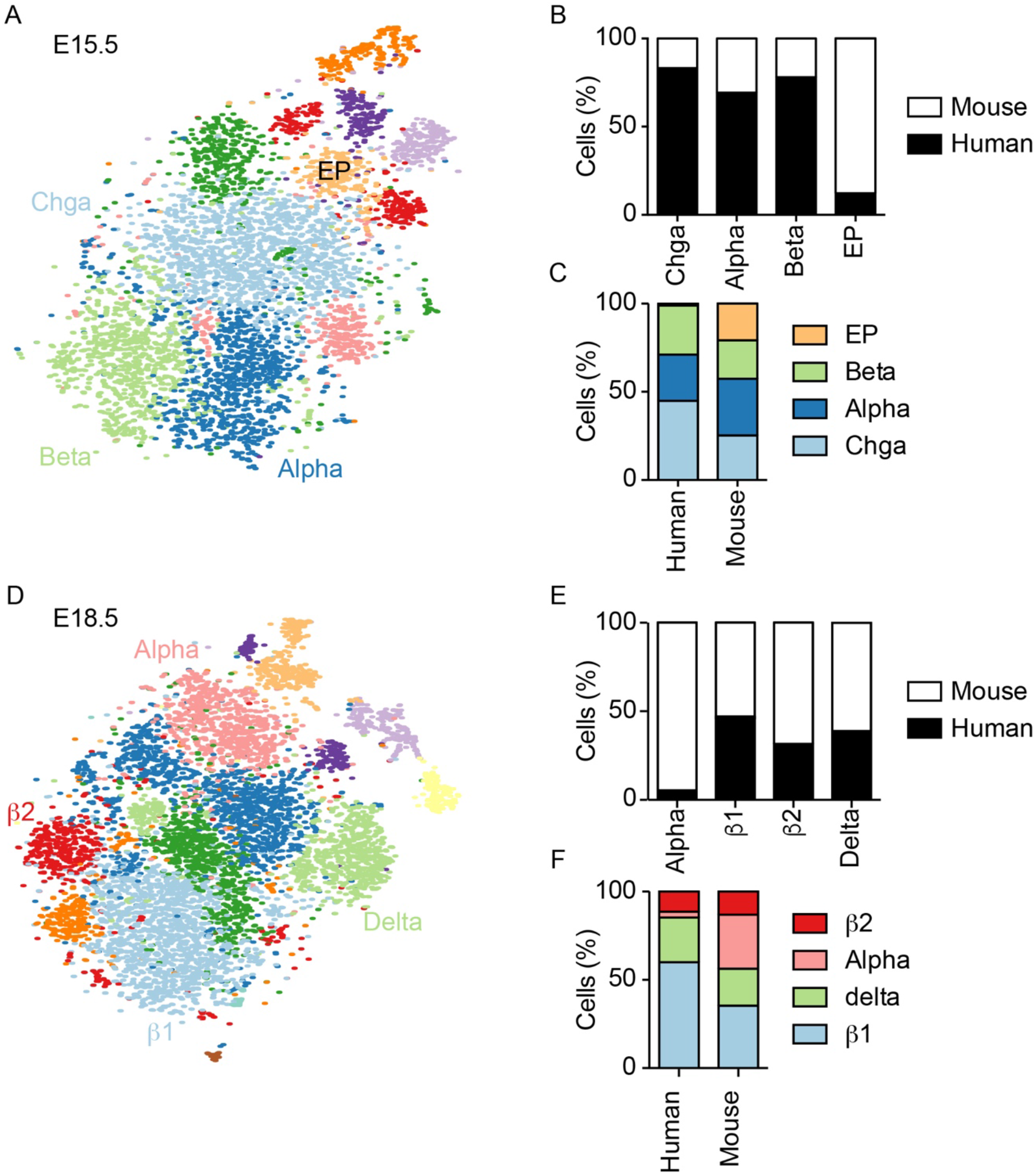
Comparison of hESC-derived endocrine cells to mouse E15.5 and E18.5 endocrine cells. (A) tSNE plot of 10 cell clusters from comparison of E15.5 yellow & green and S6D1 libraries. (B) Proportion (%) of Chga, Alpha, Beta and EP clusters that are from the mouse (white) and human (black) libraries. (C) Of the total human and mouse populations, the percentage of cells that are classified as EP, Beta, Alpha and Chga phenotype. (D) tSNE plot of 13 cell clusters from comparison of E18.5 green and S6D1 libraries. (E) Proportion (%) of Alpha, β1, β2, and Delta clusters that are from the mouse (white) and human (black) libraries. (F) Of the total human and mouse populations, the percentages of cell that are classified as β2, Alpha, Delta, and β1 phenotype.

To investigate whether the hESC-derived cells are similar to later developing endocrine cell types, the single cell transcriptome was compared to E18.5 green endocrine cells. To do this, all cells of the E18.5 green library (Figure 4A) and hESC library (Figure 6A) were merged into one dataset, unsupervised clustering was performed, and visualized using a tSNE plot revealing 13 clusters (Figure 7D). Using the endocrine cell identities of E18.5 green cells (Figure 6A), four clusters were identified as alpha cells, delta cells, immature β-cells (β2) and mature β-cells (β1). Human cells made up 47%, 39%, 32%, and 5% of the β1, delta, β2, and alpha cell clusters, respectively (Figure 7E). Of the human cells, most cells were part of the β1 and delta clusters, suggesting that that hESC-derived cells are most similar to the E18.5 mature β- and delta cells (Figure 7F).

## Discussion

Single cell RNA-sequencing allows for the identification of novel cell types, discovery of cell state specific genes, and the appreciation of cellular heterogeneity within a population. Here, scRNA-seq was used to generate a resource of single cell transcriptomes from 7,223 E15.5 embryonic pancreatic cells, 6,852 E18.5 embryonic pancreatic cells, and 4,497 hESC-derived *NEUROG3-2A-eGFP* cells. Several unique observations were made including the presence of macrophage cells during embryonic pancreas development, novel genes that may regulate endocrine cell formation, and previously unidentified populations of cells generated in hESC differentiations. This data is publically available online (https://lynnlab.shinyapps.io/embryonic_pancreas) and will serve as a resource to quickly determine the single cell expression of a particular gene in embryonic pancreas.

There were 175 and 122 macrophage cells in E15.5 and E18.5 embryonic pancreas, respectively. Macrophages are found in many tissues during development and play important roles in tissue remodeling (41). During mammary gland development, F4/80+ macrophage cells are found localized to the highly proliferative epithelial structures and are required for proper gland development (42). Using a genetic mouse model that is deficient in macrophages (*Csf1^op/op^*), macrophages were shown to play an important role in the formation of the epithelial tree during mammary gland development, a defect that can be rescued by restoring the macrophage population. The role of macrophages in the growth of epithelial organs appears to be mostly indirect, either by facilitating the clearance of apoptotic cells (43) or remodeling of extracellular matrices (44). In mouse pancreas development, macrophages are present as early as E12.5 (45, 46).

However, the presence of macrophages in the Neurog3 endocrine lineage is a novel finding. This may represent a previously unidentified developmental source of macrophages or, more likely, it results from the phagocytosis of nascent endocrine cells by macrophages, which is a previously unappreciated occurrence. Whatever the developmental source, macrophages play an important role in β-cell maturation as *Csf1^op/op^* mice have reduced β-cell mass due to decreased β-cell proliferation in late embryogenesis (46). Consistently, treating pancreatic explants with M-CSF increased the number of insulin+ cells, which is thought to be via the differentiation/activation of macrophage precursors (45). Taken together, this study confirms the presence of macrophages in embryonic pancreas and is consistent with a phagocytic role of macrophages during embryonic development.

Comparison of E15.5 and E18.5 endocrine progenitor cells resulted in a list of genes that are upregulated in endocrine progenitors. These include *Btg2* and *Gadd45a*, both of which are involved in cell cycle regulation and have been implicated in neural development. Gadd45 genes are involved in tissue development via their role in cell cycle exit and DNA demethylation (47). Pro-neural proteins, such as Neurog2, NeuroD, and Ascl1, have been shown to activate expression of *Gadd45g* (48, 49). This leads to Gadd45-dependent cell cycle exit by upregulation of cell cycle inhibitor *Cdkn1a* (50) and direct interaction with Cdk1/CyclinB (51), as has been shown in *Xenopus* embryos for both Gadd45a and Gadd45g (52). In addition, studies in mice implicate Gadd45b in reducing proliferation of neural precursors and DNA demethylation of promoters involved in adult neurogenesis (53). While Gadd45 proteins are implicated in pancreatic cancer (50), their role in pancreas development has not been investigated. The expression of *Gadd45a* in Neurog3+ endocrine progenitor cells suggest that it may play a role, along with Cdkn1a, in regulating cell cycle exit. Future studies will explore the potential role of Gadd45a in DNA demethylation of endocrine-specific promoters during pancreatic endocrine genesis

*Btg2*, also known as *Tis21*, is a negative regulator of the cell cycle that inhibits transcription of *CyclinD1*, preventing the G1-S transition (54, 55). Deletion of *Btg2* in the adult dentate gyrus shortens G1 length in progenitor cells and prevents their terminal differentiation (56). This is thought to be caused in part by the direct binding of Btg2 to *Id3* promoter. Id proteins bind E proteins, which are obligate heterodimerization partners of bHLH TFs like Neurog3. By sequestering E proteins and preventing their association with pro-neural bHLH TFs, Id acts to prevent terminal differentiation (57). It will be interesting to investigate the role of Btg2 in pancreas development. Based on literature from neurogenesis, Btg2 may also act to inhibit *Id3* transcription, allowing for the activation of pro-endocrine genes, including *Neurog3*.

Understanding β-cell maturation, a process that is thought to occur postnatally, is a key goal in regenerative medicine approaches for diabetes treatment. Postnatal β-cells are functionally immature due to a high basal insulin secretion and reduced glucose stimulated insulin secretion (58–60). This is due to reduced expression of key metabolic genes and a reduced sensitivity of ATP-sensitive K+ channel to glucose (59, 61). Recently, a single cell transcriptome study of postnatal β-cells identified an immature, proliferating phenotype that was marked by high expression of mitochondrial and amino acid metabolism genes (3). Studies on both rat and human postnatal β-cells suggest that maturation of β-cell function, i.e. glucose stimulated insulin release, occurs postnatally (62, 63) and this is concomitant with a decline in β-cell proliferation. Furthermore, when adult β-cells are induced to proliferate, they resemble functionally immature neonatal cells (30), suggesting that proliferation of β-cells leads to a decline in function. However, in embryonic proliferating β-cells we did not detect significant changes in gene expression that suggests a defect in function, likely owing to the immature state of embryonic β-cells. Future studies investigating the single cell transcriptome of maturing, postnatal β-cells may provide insights into how β-cell function develops.

The liver, like the pancreas, is derived from the foregut endoderm. The region of the endoderm that gives rise to the liver can also form the ventral pancreas (64). One of the factors that controls the decision between liver and pancreas is the secretion of FGFs by the cardiac mesoderm that permits the formation of liver, while preventing ventral pancreas formation (65). The similar developmental origin of the pancreas and the liver makes the unintended generation of liver cells during hESC differentiations towards pancreas likely. However, finding liver cells downstream of *NEUROG3* is surprising. It is possible that, like in the mouse, a small population of hESC-derived pancreatic endoderm cells have low transcription of *NEUROG3* that is not sufficient to induce the endocrine lineage. Using a *NEUROG3* lineage tracing hESC line, it would be interesting to investigate the plasticity of cells that activate *NEUROG3* transcription. Our previous studies suggest that NEUROG3 protein in hESC differentiations is hyperphosphorylated (Nicole Krentz and Francis Lynn unpublished results), likely resulting in rapid degradation. Efforts to stabilize NEUROG3 protein may prevent the unintended formation of other endodermal cell types, including liver cells.

In conclusion, the single cell transcriptome of mouse pancreatic progenitors, endocrine progenitors, and endocrine cells at E15.5 and E18.5 as well as *NEUROG3*-expressing cells derived from hESCs was characterized. These data are a resource for developmental biologists interested in studying heterogeneity in the developing mouse pancreas and for stem cell researchers aiming to improve the current differentiation protocols for generating β-like cells.

## Experimental Procedures

### Animals

Mice were housed on a 12-hour light-dark cycle in a climate-controlled environment according to protocols approved by the University of British Columbia Animal Care Committee. *Rosa26^mT/mG^* (Stock No: 007576) (66) and *Neurog3-Cre* (Stock No: 005667) mice were purchased from Jackson Laboratory.

### Maintenance and in vitro differentiation of pluripotent cells

Previously, a *NEUROG3-2A-eGFP* (N5-5) reporter hESC line was generated (32) from CyT49 parental hESC line (ViaCyte, Inc. San Diego CA). Undifferentiated cells were maintained on diluted Geltrex-coated (ThermoFisher Scientific; 1:100 in DMEM/F12) plates in 10/10 media [DMEM/F12, 10% XenoFree KnockOut™ Serum Replacement (ThermoFisher Scientific), 1x MEM non-essential amino acids (ThermoFisher Scientific), 1x Glutamax, 1x penicillin/streptomycin, 10 nM β-mercaptoethanol (Sigma-Aldrich), supplemented with 10 ng/mL ACTIVIN A (E-biosciences) and 10 ng/mL HEREGULIN-β1 (Peprotech)] (67, 68). Cells were split every three or four days and plated at a density of 1×10^6^ per 60 mm plate.

For differentiation, N5-5 hESCs were plated onto Geltrex-coated 12-well plates at a density of 5 x10^5^ in 10/10 media. Differentiations began 48 hours post-seeding using a modified version of Rezania *et al* (33). Briefly, cells were rinsed with 1 x PBS and then basal culture media (MCDB 131 medium (USBiological Life Sciences), 1.5 g/L sodium bicarbonate (Sigma-Aldrich), 1 x Glutamax (ThermoFisher Scientific), 1 x P/S (ThermoFisher Scientific)) with 10 mM final glucose (Sigma-Aldrich), 0.5% BSA (Sigma-Aldrich), 100 ng/mL ACTIVIN A, and 3 μM of CHIR-99021 (Sigma-Aldrich) was added for 1 day only. For the following two days, cells were treated with the same media without CHIR-99021 compound to generate definitive endoderm (Stage 1). On day four, cells were cultured in basal media with 0.5% BSA, 10 mM glucose, 0.25 mM ascorbate (Sigma-Aldrich) and 50 ng/mL of KGF (R&D or StemCell Technologies) for 2 days to generate primitive gut tube (Stage 2). To produce posterior foregut (Stage 3), cells were treated for three days with basal media with 10 mM final glucose concentration, 2% BSA, 0.25 mM ascorbate, 50 ng/mL of KGF, 0.25 μM SANT-1 (Tocris Biosciences), 1 μM retinoic acid (Sigma-Aldrich), 100 nM LDN193189 (EMD Millipore), 1:200 ITS-X, and 200 nM α-Amyloid Precursor Protein Modulator (APPM; EMD Millipore). For stage 4, cells were treated with basal media with 10 mM glucose, 2% BSA, 0.25 mM ascorbic acid, 2 ng/mL of KGF, 0.25 μM SANT-1, 0.1 μM retinoic acid, 200 nM LDN193189, 1:200 ITS-X, and 100 nM APPM for 3 days to generate pancreatic progenitors. Cells were maintained as planar cultures and media was changed to basal media with 20 mM glucose, 2% BSA, 0.25 μM SANT-1, 0.05 μM retinoic acid, 100 nM LDN193189, 1:200 ITS-X, 1 μM T3 (Sigma-Aldrich), 10 μM Repsox (Sigma-Aldrich), and 10 μM zinc sulfate (Sigma-Aldrich) for 3 days to generate pancreatic endocrine precursors (Stage 5).

### Preparing cells for single cell RNA-sequencing

For mouse studies, *Neurog3-Cre*; *Rosa26^mTmG^* embryos were collected on E15.5 and E18.5 and dissected on ice. To generate single cells, embryonic pancreases were incubated in 2 mL of pre-warmed 37°C 0.25% Trypsin with mild agitation for 8 or 20 minutes for E15.5 and E18.5 pancreases, respectively. To stop digestion, 1 mL of cold FBS and 2 mL of cold PBS were added and mixed by inversion to stop digestion, followed by pipette filtering with a 40 μm nylon filter. Cells were then centrifuged at 4°C for 5 minutes at 200 xg. After aspirating the supernatant, cells were resuspended in cold 2% FBS in PBS, placed on ice, and immediately sorted into tdTomato+, tdTomato+eGFP+ (yellow), and eGFP+ fractions using a Beckman Coulter MoFlo Astrios (Mississauga, ON, Canada) into 20% FBS in PBS.

For hESC studies, N5-5 cells were differentiated and collected following three days at stage 5 (S6D1). Cells were washed once with PBS before 500 μL of Accutase was added per well of a 12-well plate. Following 5 minutes at 37°C, 500 μL of 2% BSA MCDB media was added to each well and cells were transferred to a 15 mL conical tube. Cells were centrifuged for 5 minutes at 200 xg, washed once with PBS, and resuspended in 350 μL of ice cold PBS. GFP+ cells were sorted into stage 5 media with 10 μM Y-27632 dihydrochloride using a Beckman Coulter MoFlo Astrios.

### Generating scRNA-seq libraries

The 10x Genomics Chromium^TM^ controller and Single Cell 3’ Reagent Kits v2 (Pleasanton, CA, USA) were used to generate single cell libraries. Briefly, cells were counted following FACS and cell suspensions were loaded for a targeted cell recovery of 3000 cells per channel. The microfluidics platform was used to barcode single cells using Gel Bead-In-Emulsions (GEMs). RT is performed within GEMs, resulting in barcoded cDNA from single cells. The full length, barcoded cDNA is PCR amplified followed by enzymatic fragmentation and SPRI double sided size selection for optimal cDNA size. End repair, A-tailing, Adaptor Ligation, and PCR are performed to generate the final libraries that have P5 and P7 primers compatible with Illumina sequencing. The libraries were pooled and sequenced using an Illumina NextSeq500 platform with a 150 cycle High Output v2 kit in paired-end format with 26 bp Read 1, 8 bp I5 Index, and 85 bp Read 2.

### Data Analyses

Following sequencing, data were analyzed using publically available software programs and R pipelines. First, cellranger mkfastq (10x Genomics) generates FASTQ files from the raw sequencing data, storing the nucleotide sequence and its corresponding quality score in a text-based format for further analysis. Next, cellranger count uses the FASTQ file to perform sequence alignment (mouse: GRCm38 and human: GRCh38), filter sequences based on quality score, and generate single cell gene counts. As an optional step, cellranger aggr can be used to combine data from multiple samples. This was used to merge all E15.5 and E18.5 libraries into E15.5 total cells and E18.5 total cells datasets, respectively.

As minimal filtering is performed in cellranger, two additional R pipelines were used to filter out cells that did not meet the quality control standard. The first pipeline is called Scater (https://bioconductor.org/packages/release/bioc/html/scater.html) and is a single cell analysis pipeline that places a great emphasis on quality control (69). Scater discards cells based on the total number of expressed genes, removing potential doublets and debris, and removes low-abundance genes or genes with high dropout rate based on expression level. For this analysis, cells were discarded based on counts (transcripts/gene) or genes (genes/cell) greater than 3 standard deviation away from the mean. This QC dataset was then analyzed using the Seurat V2.0 pipeline (http://satijalab.org/seurat/), another R toolkit for single cell genomics (70). Seurat was used to remove common sources of variation including number of genes (each cell must express a minimum of 500 genes), number of counts (each gene must be expressed in a minimum of three cells), and cell cycle phase. Finally, unsupervised *k*-means clustering was performed using Seurat to group cells based on gene expression and to identify unique cell types within the populations.

### Identification of cell cycle phase

To identify the cell cycle stage of individual cells, each cell was assigned a score based on expression of G2/M and S phase markers using Seurat v2.0. From this score, a cell is classified as either G1-phase (expressing neither G2/M or S phase markers), S-phase (expressing only S markers), or G2/M-phase (expressing only G2/M markers). This data can then be used to regress out the cell cycle phase as a source of heterogeneity.

### Pseudotime analysis

Pseudotime analysis was performed using Monocle v2.6.1 (http://cole-trapnell-lab.github.io/monocle-release/docs/#constructing-single-cell-trajectories). Transcript data that was quality controlled using Scater was loaded into Monocle as a CellDataSet object. Variable expressed genes was defined as a gene that was expressed in >50 cells. Unsupervised clustering was performed using genes that have a mean expression of ≥ 0.1 and dimensional reduction was done using tSNE. Next, differential gene expression analysis was done between clusters of interest and the top 1000 variable genes were used to order cells in the pseudotime.

## Author Contributions

Conceptualization, N.A.J.K., E.E.X, S.S. and F.C.L.; Methodology and Investigation, N.A.J.K., E.E.X., M.L., S.S., and F.C.L. Writing, N.A.J.K. and F.C.L.; Funding Acquisition, F.C.L.

## Acknowledgements

This work was supported by operating grants to F.C.L.: Stem Cell Network (DT3 and DT4), Canadian Foundation for Innovation (#33644). Salary (F.C.L.) was supported by the Michael Smith Foundation for Health Research (#5238 BIOM), the Canadian Diabetes Association, and the BC Children’s Hospital Research Institute. Fellowship support was provided by the CIHR-BC Transplantation Trainee Program, the BC Children’s Hospital Research Institute, UBC, and the National Science and Engineering Research Council of Canada PGSD2-475838 (N.A.J.K.). We thank members of the Lynn lab for technical support, discussion, and critical reading of the manuscript. Training on the 10X genomics platform was provided by Jens Durruthy (10X Genomics).

## Supplemental Information titles and legends

**Figure S1: Top ten differentially expressed genes in trunk and endocrine cell populations at E15.5 and E18.5, related to Figure 1.**

(A) Single cell expression of top ten differentially expressed genes in trunk (green), endocrine progenitors (EP; light green), and endocrine cells (yellow) at E15.5. (B) Single cell expression of top ten differentially expressed genes in trunk (green) and two immature endocrine clusters (blue and teal) at E18.5.

**Figure S2: Characterization of mouse E15.5 yellow and green cells, related to Figure 2.**

(A) tSNE plot identifying cell cycle phase of individual cells in E15.5 yellow and green population. G1-phase in red, S-phase in blue and G2/M-phase in green. (B) Single cell gene expression of *Neurog3*, *Neurod1*, *Gcg*, *Ghrl*, *Ins1*, and *Ins2*. (C) Heatmap of top ten differentially expressed genes in Chga (green), trunk (blue), Ghrl (purple), and Macrophage (pink) clusters. (D) Single cell gene expression of *tdTomato*, *eGFP* and trunk cluster specific genes (*Spp1*, *Cxcl12*, *Cyr61*, *Mt1*, *Mt2*, *Cpa1*, and *Cpa2*).

**Figure S3: Analysis of lineage specified trunk progenitor cells at E15.5, related to Figure 3.**

(A) Heatmap of top ten genes expressed in the trunk cells along the acinar, ductal and endocrine lineage.

**Figure S4: Differentially expressed genes in E18.5 endocrine cell clusters, related to Figure 4.**

(A) Heatmap of top ten genes expressed in β1 (red), β2 (green), β3 (blue), and S-phase (purple) clusters in E18.5 green cells. (B) Heatmap of top ten genes expressed in trunk (green) and endocrine progenitor (EP; pink) clusters in E18.5 green cells.

**Figure S5: Differentially expressed genes in E18.5 green non-β-cell endocrine clusters, related to Figure 5.**

(A) tSNE plot of individual cell cycle phase of E18.5 green cells. (B) Single cell gene expression of endocrine hormones *Ins1*, *Ins2*, *Ppy*, *Gcg*, *Sst*, and *Ghrl*. (C) Heatmap of top ten differentially expressed genes in α-cells (orange), δ-cells (yellow), PP-cells (green), and Ghrl cells (blue).

## Supplementary Tables

**Table S1: Differential Expression Analysis of E15.5 mouse pancreatic cells.** Related to Figure 1. Lists of genes that are differentially expressed in each cluster of E15.5 total pancreatic cells. Within the excel file, each cluster has its own sheet where the differentially expressed genes are listed in descending order by average differential expression and includes relevant p value.

**Table S2: Differential Expression Analysis of E18.5 mouse pancreatic cells.** Related to Figure 1. Differentially expressed gene lists for each cluster of E18.5 total pancreatic cells. The differentially expressed genes per cluster (individual sheets within excel file) are listed in descending order based on average differential expression.

**Table S3: Differential Expression Analysis of E15.5 endocrine-lineage cells.** Related to Figure 2. Differentially expressed gene lists for each cluster of E15.5 yelow and green cells. The differentially expressed genes per cluster (individual sheets within excel file) are listed in descending order based on average differential expression.

**Table S4: Differential Expression Analysis of E18.5 endocrine-lineage cells.** Related to Figure 4. Lists of genes that are differentially expressed in each cluster of E18.5 yellow and green cells. Within the excel file, each cluster has its own sheet where the differentially expressed genes are listed in descending order by average differential expression and includes relevant p value.

**Table S5: Differential Expression Analysis of E18.5 endocrine cells.** Related to Figure 5. Lists of genes that are differentially expressed in each cluster of E18.5 green cells. Within the excel file, each cluster has its own sheet where the differentially expressed genes are listed in descending order by average differential expression and includes relevant p value.

