## Supplementary Materials for "Single cell transcriptome profiling of mouse and hESC-derived pancreatic progenitors"

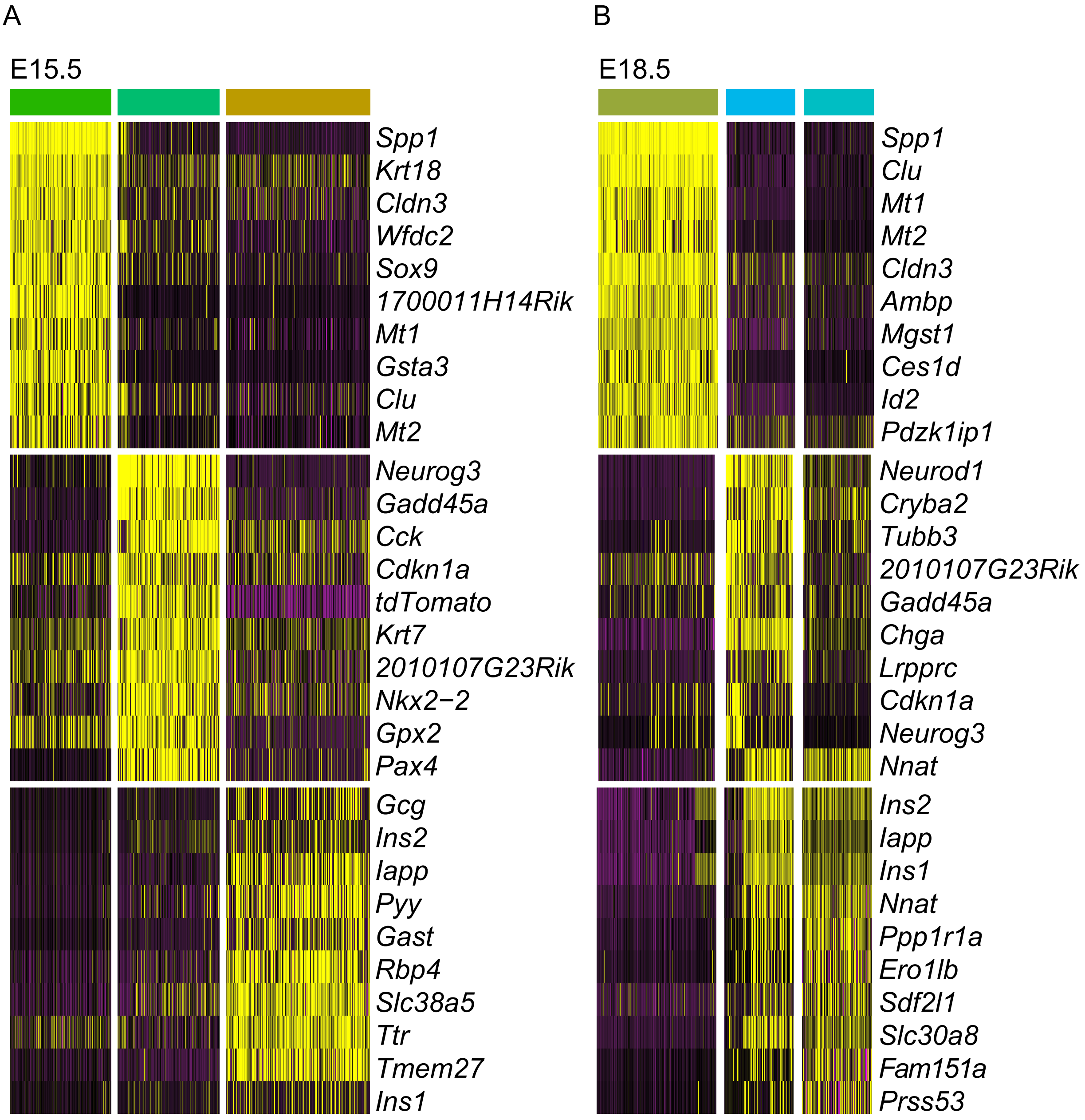


**Figure S1: Top ten differentially expressed genes in trunk and endocrine cell populations at E15.5 and E18.5, related to Figure 1.**
(A) Single cell expression of top ten differentially expressed genes in trunk (green), endocrine progenitors (EP; light green), and endocrine cells (yellow) at E15.5 (C) Single cell expression of top ten differentially expressed genes in trunk (green) and two immature endocrine clusters (blue and teal) at E18.5.

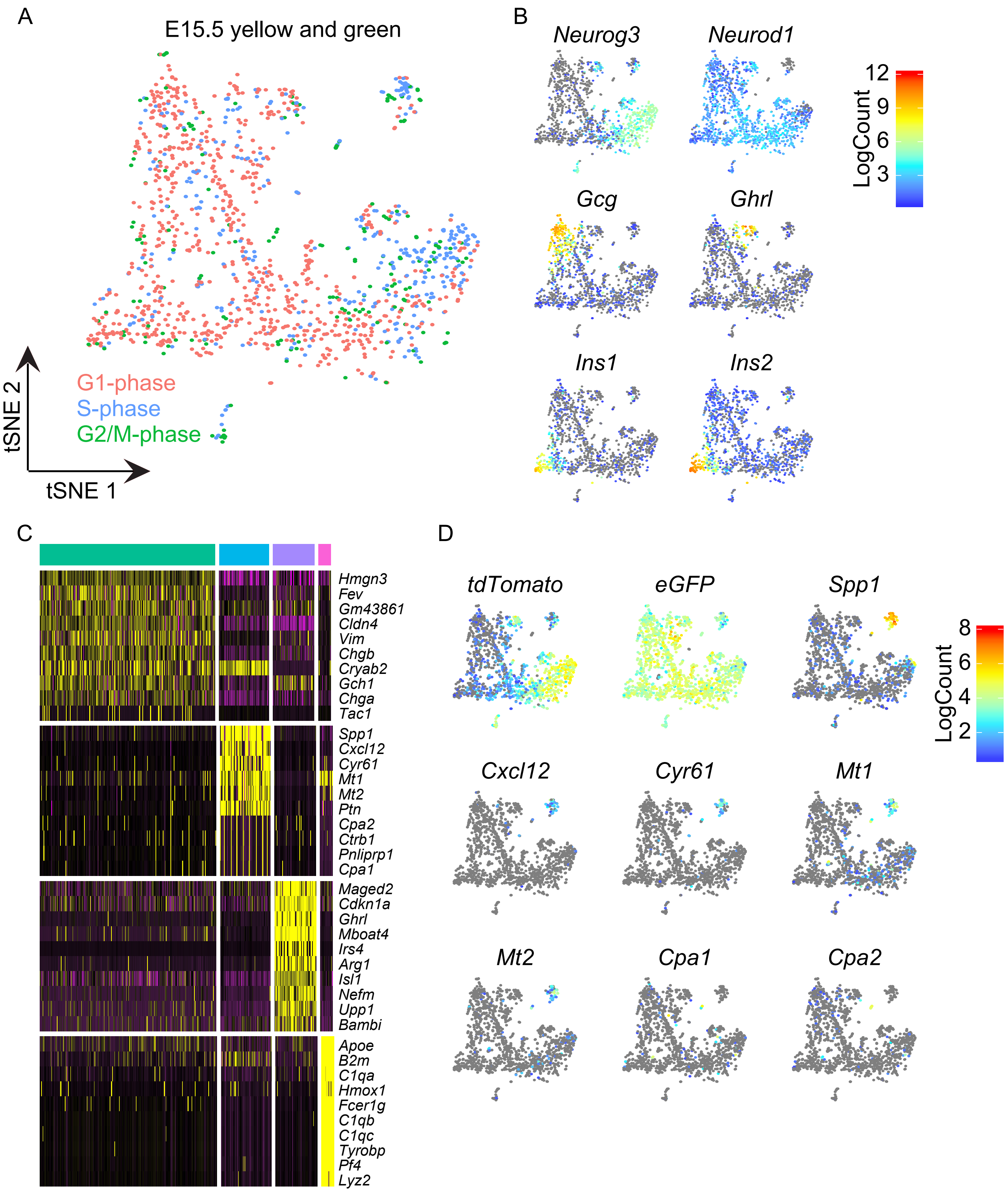


**Figure S2: Characterization of mouse E15.5 yellow and green cells, related to Figure 2.**
(A) tSNE plot identifying cell cycle phase of individual cells in E15.5 yellow and green population. G1-phase in red, S-phase in blue and G2/M-phase in green. (B) Single cell gene expression of *Neurog3*, *Neurod1*, *Gcg*, *Ghrl*, *Ins1* and *Ins2*. (C) Heatmap of top ten differentially expressed genes in Chga (green), trunk (blue), Ghrl (purple), and macrophage (pink) clusters. (D) Single cell gene expression of *tdTomato*, *eGFP*, and trunk cluster specific genes (*Spp1*, *Cxcl12*, *Cyr61*, *Mt1*, *Mt2*, *Cpa1*, and *Cpa2*).

A

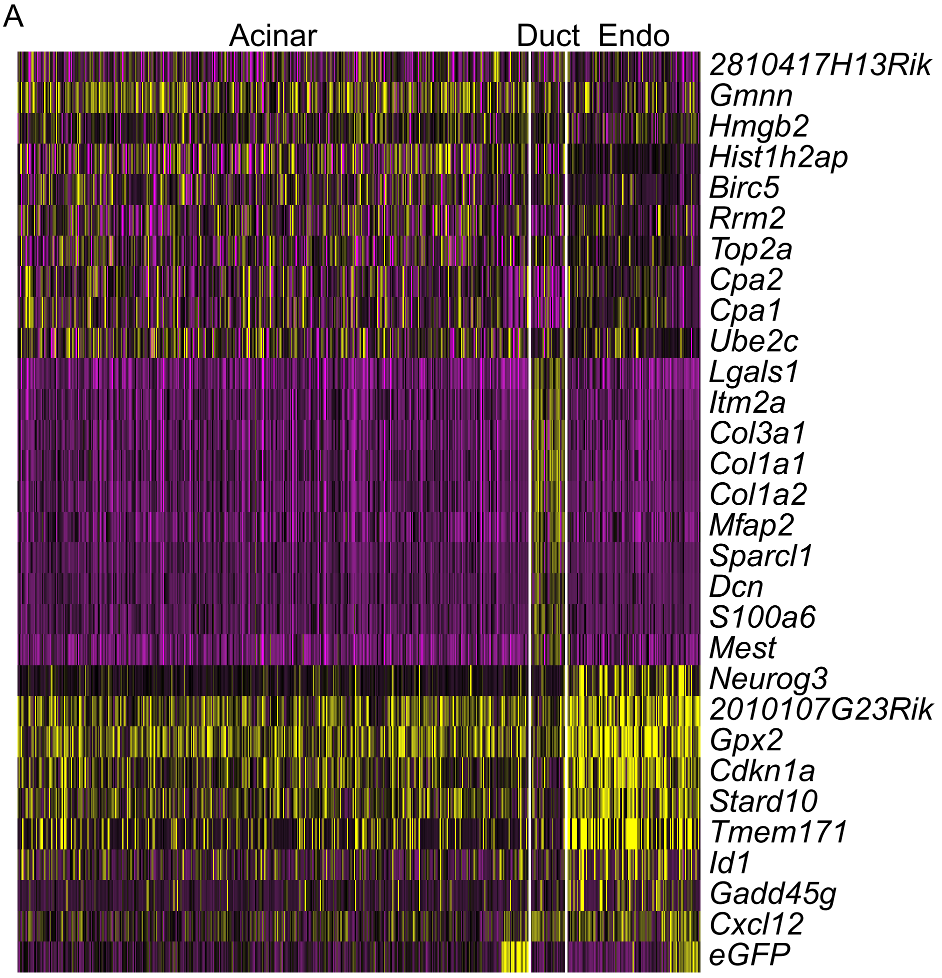


**Figure S3: Analysis of lineage specified trunk progenitor cells at E15.5, related to Figure 3.**
(A) Heatmap of top ten genes expressed in the trunk cells along the acinar, ductal, and endocrine lineage.

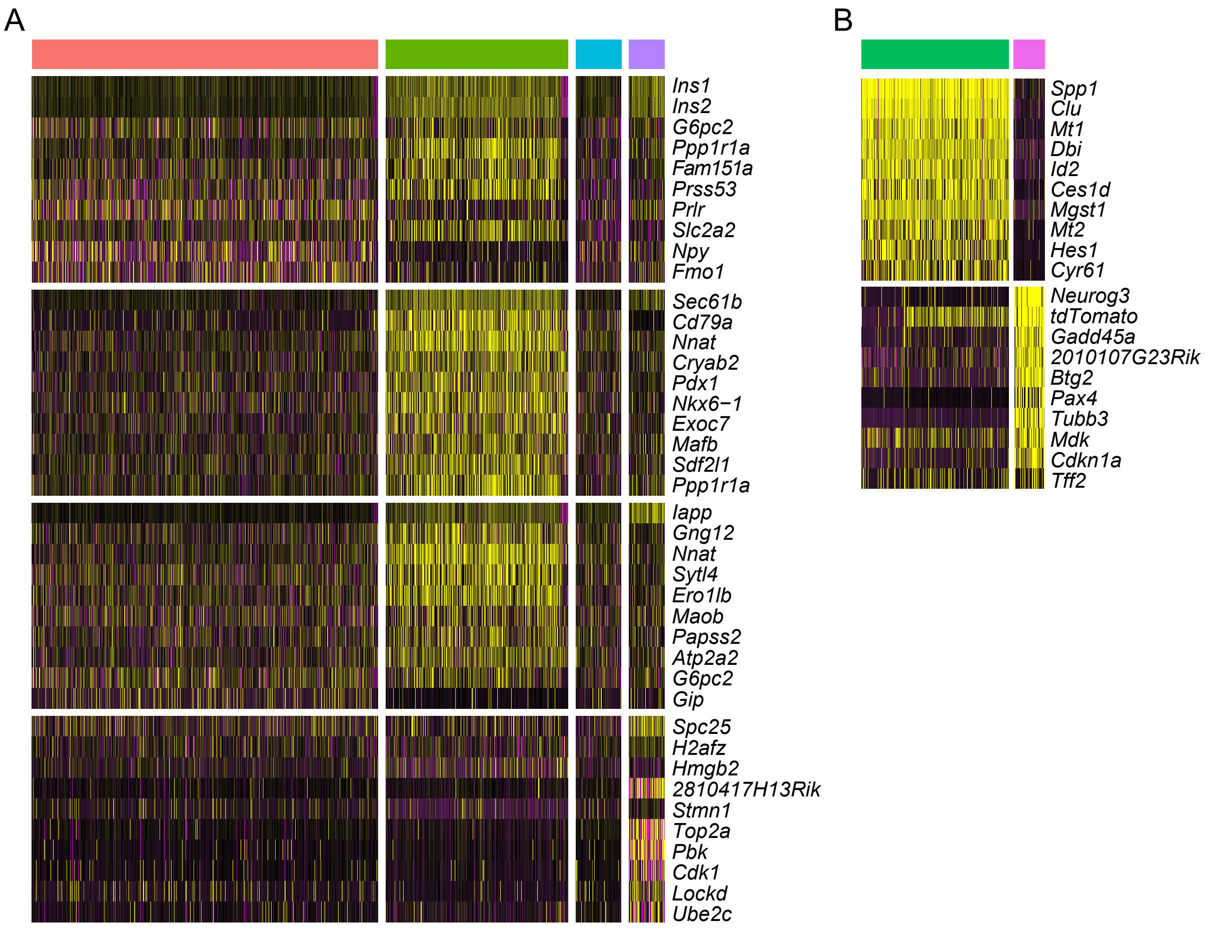


**Figure S4: Differentially expressed genes in E18.5 endocrine cell clusters, related to Figure 4.**
(A) Heatmap of top ten genes expressed in β1 (red), β2 (green), β3 (blue), and S-phase (purple) clusters in E18.5 green cells. (B) Heatmap of top ten genes expressed in trunk (green) and endocrine progenitors (EP; pink) clusters in E18.5 green cells.

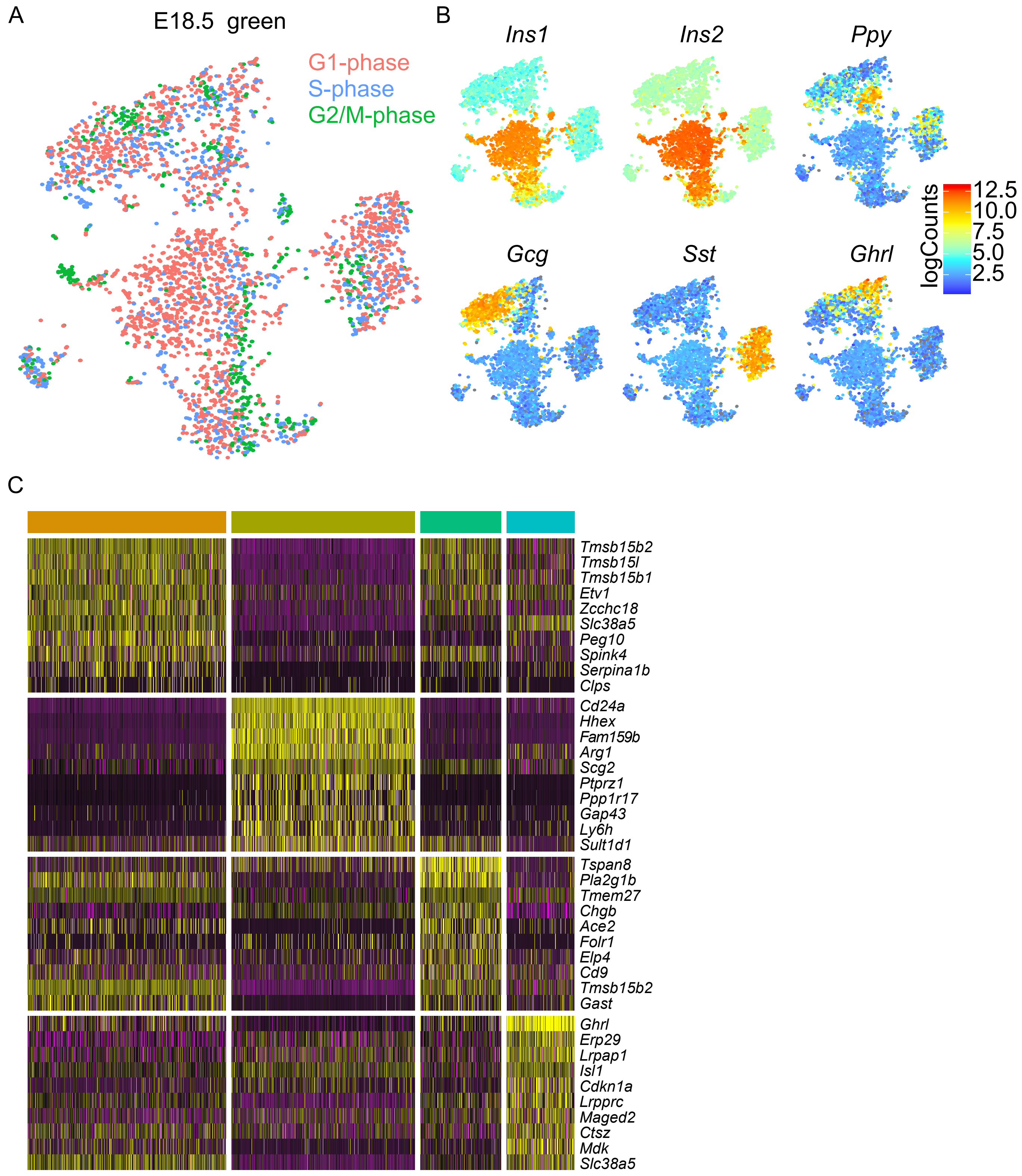


**Figure S5: Differentially expressed genes in E18.5 green non-β-cell endocrine clusters, related to Figure 5.**
(A) tSNE plot of individual cell cycle phase of E18.5 green cells. (B) Single cell gene expression of endocrine hormones *Ins1*, *Ins2*, *Ppy*, *Gcg*, *Sst*, and *Ghrl*. (C) Heatmap of top ten differentially expressed genes in α-cells (orange), δ-cells (yellow), PP-cells (green), and Ghrl cells (blue).
